# Using Deep Learning with Different Architectures to Recognize RNA:DNA Triplex Structures from Histone Modification Features

**DOI:** 10.1101/2025.09.16.676231

**Authors:** Joseph L. Tsenum

## Abstract

Long non-coding RNAs (lncRNAs) can perform their regulatory roles by forming triple helices through RNA-DNA interaction. Although this has been verified by few in vivo and in vitro methods, in silico approaches that seek to predict the potentials of lncRNAs and DNA sites becoming a triplex forming structure is required. Triplexator have also predicted vast amounts of lncRNAs and DNA sites that has the potentials of becoming a triplex structure. There is also an emerging experimental-evidence that the presence of epigenetic marks at DNA sites and lncRNAs can facilitate the formation of RNA:DNA triplex structures. There is therefore, a huge demand for computational approaches such as deep learning that can make novel predictions about RNA:DNA triplex structure formation. In this study, we developed four (4) deep neural network models that can predict the potentials of lncRNAs and DNA sites to form triple helices genome-wide, by taking histone modification marks as features. Our data was first passed through the Triplexator to screen out lncRNAs and DNA sites with low potentials of forming triple helices. We used different deep learning architectures to build our models, including two-layer convolutional neural networks (CNN) and multilayer perceptron (MLP). Our DNA2_CNN model performed best at a mean AUC of 0.78 at 32 Kernel size and learning rate of 1e-3. Our deep neural network models revealed several novel lncRNAs and DNA sites, including HOTAIR, MEG3, PARTICLE, DACOR1, MIR100HG, FENDRR, ANRIL, TUG1, MALAT1, LINC00599, TINCR, NEAT1, roX2, DHFR, OTX2-AS1, Xist, SNHG16, ATXN8OS, BCYRN1, TERC, Khps1, that have the potential of forming triplex structures, thereby confirming previous experimental results and that of the Triplexator. The performance of our models also supports previous findings that histone modification marks can help in identifying lncRNAs and DNA regions that have the potentials of forming RNA:DNA triplex structures. In conclusion, we showed that different deep learning architectures can recognize lncRNAs and DNA that have the potentials of forming RNA:DNA triplex structures.

## Introduction

Advances in genome sequencing technologies in the past decade have shown the complexity of genome organization in both eukaryotic and prokaryotic organisms (Kapranov *et al*., 2007; ENCODE Project Consortium, 2004, 2012). These genome sequencing technologies has revealed large amounts of non-coding RNAs (ncRNAs) that were previously coined as ‘junk DNA’ and a small fraction of protein-coding mRNA in the mammalian DNA.

Efforts have been made in the past years in sequencing ncRNAs and quantifying their various regulatory roles (Quinn and Chang, 2016; Bonasio and Shiekhat-tar, 2014; Batista and Chang, 2013; Lee, 2012). Non coding RNAs (ncRNAs) consists of a broad class of regulatory RNAs; these include small ncRNAs such as small interferingRNAs (siRNAs), PIWI-interacting RNAs, microRNAs (miRNAs), and long noncoding RNAs (lncRNAs), which are made up of more than 200 nucleotide in length (Cech and Steitz, 2014; Kowalczyk *et al*., 2012). Majority of the mammalian non-coding RNAs are highly conserved, suggesting that they perform significant functions in biological processes (Struhl, 2007).

Long non-coding RNAs (lncRNAs) interacts with epigenetic factors to structure the results of important biological roles such as gene transcription regulation, modulation of nuclear organization, pre-mRNA splicing and X inactivation in females. This has improved our knowledge of the interaction between lncRNAs and epigenetic factors of human diseases as biomarkers of genetic diseases and their therapeutic targets (David J. H *et al*., 2018).

Long non-coding RNAs (lncRNAs) are capable of forming RNA-DNA hybrid duplexes and RNA-DNA triplexes structures at specific DNA sequences (Wang and Chang, 2011; Li *et al*., 2016). These triplex-forming lncRNAs are important in biological processes. Both *in-vivo* and *in-vitro* methods have verified the potential of some lncRNAs in forming triplex structures with DNA sequences. Various mathematical, statistical and computational methods have also been applied in predicting lncRNAs that have the potential of forming triplex structures with specific DNA regions. Computationally, classical machine learning and deep neural networks have shown promise in predicting these triplex-forming lncRNAs and DNA sites.

## Aims of the Research

Research on the functions of long non-coding RNA as decoys, scaffolds, signals, and guides, to function in gene expression regulation and chromatin states modulation is emerging rapidly (Mercer and Mattick, 2013). Recent sequencing technologies such as CHART-Seq, CHOP-Seq, ChIRP-Seq and RAP-Seq has led to the discovery of large amounts of long non-coding RNAs (lncRNAs). Recent research has shown that long non-coding RNAs (lncRNAs) can form triplex structures with the DNA to perform their regulatory roles (Zhang, *et al*., 2020). They form these structures by interacting with the DNA, other RNAs and proteins in order to guide regulatory proteins, localizing to specific loci thereby shaping the three-dimensional (3D) nuclear organization (Mercer and Mattick, 2013). Previous research that described the canonical B-form DNA double helix also proposed that other possible DNA, DNA-RNA, and RNA structures that plays vital roles as functional genomic elements exists (Albino *et. al*., 2015). Mathematical and statistical base pairing rules such as Triplexator, Triplex Domain Finder (TDF), LongTarget and Triplex-Inspector have been applied over the years in predicting DNA:RNA triplex structure formation (Zhang *et al*., 2020). These methods have been known to recognize vast amounts of potential triplex forming lncRNAs and DNA sites based on canonical rules (Zhang *et al*., 2020). Little amounts of experimentally verified triplex-forming lncRNAs suggests that some of the predicted triplex-forming lncRNAs by canonical rules may not form triplex structures in practice (Zhang *et al*., 2020), however more research is needed to predict the exact triplex-forming lncRNAs and DNA sites. Theoretical triplex potential can be done by computational methods without taking into account *in vitro* and *in vitro* assays verified data.

There is therefore an increasing demand for computational methods such as deep neural networks that can predict triplex-forming lncRNAs and DNA sites. Deep learning has been successfully applied on omics studies in recent years. In recent years, it has been established that convolutional neural networks achieve higher performance in predicting triplex-forming lncRNAsand DNA sites using DNA sequences than classical machine learning methods. However, low performance predictions may suggest that the information stored in the primary DNA sequence is not sufficient for building models effectively. In the present study therefore, we aim at recognizing these triplex-forming lncRNAs and DNA sites by deep learning architectures by taking histone modification marks as features. We would add histone modification marks to our DNA sequences to ascertain if there will be improvement from low quality predictions as reported in other previous works. We intend to use different deep learning architectures such as convolutional neural networks (CNN), and multilayer perceptron (MLP) and compare the performance of our models to know the best performing deep learning architecture in predicting triplex-forming lncRNAs and DNA sites.

## Background

### Non-Coding RNA and its Types

Sequencing of the human genome became a breakthrough in our knowledge of the major role of RNAs in gene expression regulation and chromatin modulation. Out of the sequenced human genome, 2% accounted for protein-coding genes while 98% accounted for non-coding genes, in spite of more than 90% of the genome being transcribed (Wilusz *et al*., 2009; Birney *et al*., 2007; Katayama *et al*., 2005). This was contrary to earlier believes on how cells of the mammalian origin work. Further research has revealed the roles of non-coding RNAs like lncRNAs in genome regulation and the etiology of human diseases (Maass *et al*., 2014; Rinn *et al*., 2007; Costa 2005). Non coding RNAs such as small ncRNAs (small interfering RNAs (siRNAs), PIWI-interacting RNAs, microRNAs (miRNAs), small nucleolar RNAs (snoRNAs) (Taft *et al*., 2010)); and long noncoding RNAs (lncRNAs), which are made up of more than 200 nucleotide in length were recently profiled as key regulators of gene regulation and chromatin modulation. The recent finding of an interesting group of regulators known as long non-coding RNAs (lncRNAs) has given us novel understanding in biology to explore for the etiology of different diseases ranging from cancer sub-typing, sepsis, aging, cardiovascular diseases to other human diseases. This knowledge has helped in recent times for therapeutic interventions of different ailments. Today, lncRNAs are known to be key classes of the varying RNA family in the living cells as shown in Figure 2.1A. Most definitions describing lncRNAs and their analogue transcripts can be found in Figure 2.1B.

They primarily regulate gene expression and epigenetic control of chromatin in many different cellular systems and pathways, X-chromosome inactivation and imprinting, promoter-specific gene regulation and mRNA stability (Hombach and Kretz, 2016). Mammals use Xist long non-coding RNA to silence (inactivate) the extra X chromosome in females to balance expression level in the two sexes. Xist induces a repressive epigenetic landscape thereby creating the inactiveX chromosome known as Xi (Charles and Ogawa, 2015). These long non-coding RNAs plays important roles in developmental stages of living things.

### Long Non-Coding RNA and its Functions

Recently, the discovery of important biological molecules called long noncoding RNAs (lncRNAs) has brought a new dimension of diagnostic and therapeutic opportunities for the fight against human diseases. NONCODE database is a ncRNAs database (excluding tRNAs and rRNAs), are databases that contain over 100, 000 lncRNAs, out of which majority of the lncRNAs are yet to be annotated (Fang *et al*., 2018). It has long been established that tRNAs and rRNAs have structural role in biological processes without taking part in translation (Rinn and Chang, 2012; Mattick, 2011). Long lncRNAs have more than 200 nt in length (Amaral *et al*., 2011). Long non-coding RNAs can be found in animals (Clemson *et al*., 1996; Brown *et al*., 1992), plants (Swiezewski *et al*., 2009), viruses (Reeves *et al*., 2007), yeast (Houseley *et al*., 2008), and prokaryotes (Bernstein *et al*.,1993). In comparison with other well-studied RNAs such as mRNAs, miRNAs, and snoRNAs, (Derrien *et al*., 2012; Pang *et al*., 2006; Wang *et al*., 2004), lncRNA’s inter-specie conservation is poor which explains why its functionally is questioned. Research has shown that lncRNAs usually have low expression levels (Derrien *et al*., 2012) thereby being viewed as noise during transcription. This notwithstanding, the last two decades has provided us with several researches signifying that lncRNAs acts in different biological processes (Clark *et al*., 2011, Ponting *et al*., 2009) such as transcription (Martianov *et al*., 2007; Rinn *et al*., 2007), splicing (Tripathi *et al*., 2010; Rintala-Maki and Sutherland, 2009), translation (Beltran *et al*., 2008; Muddashetty *et al*., 2002), protein localization (Campalans *et al*., 2004; Watanabe and Yamamoto, 1994), cellular structure integrity (Sunwoo *et al*., 2009; Kloc *et al*., 2005), imprinting (Bartolomei *et al*., 1991; Brockdorff *et al*., 1991; Brown *et al*., 1991), cell cycle (Mourtada-Maarabouni *et al*., 2008; Ginger *et al*., 2006), and apoptosis (Huarte *et al*., 2010; Reeves *et al*., 2007), stem cell pluripotency (Sheik *et al*., 2010), reprogramming (Loewer *et al*., 2010), and heat shock response (Shamovsky *et al*., 2006; Espinoza *et al*., 2004). Scientists have demonstrated that lncRNAs may play a vital role in regulating cancer progression (Tano and Akimitsu, 2012). The roles of lncRNAs in the development of many other human diseases was also demonstrated by Wapinski and Chang, 2011.

Large amounts of lncRNAs have been profiled by different sequencing techniques. This has given us understanding on how these lncRNAs functions as as signals, decoys, scaffolds, guides, enhancer RNAs, and short peptides (Li *et al*., 2014; Moran *et al*., 2012). As signals, lncRNAs functions as molecular signals in response to stimuli during transcriptional regulation (Pandey *et al*., 2008). Decoy lncRNAs functions to reduce the availability of regulatory factors such as transcription factors, subunits of larger chromatin modifying complexes, miRNAs, and catalytic proteins by introducing what is known as “decoy” binding sites (Kallen *et al*., 2013). As scaffolds, multiple-component complexes, such as ribonucleoprotein (RNP) complexes are assembled on structures provided by lncRNA scaffold transcripts (Yang *et al*., 2014). For ribonucleoprotein (RNP) complexes, transcriptional activation or repression can occur depending on the nature of proteins and RNAs present as soon as the complexes have been fully assembled on the “decoy” binding sites (Aguilo *et al*., 2011; Kotake *et al*., 2011). In this particular case, guide lncRNAs usually directs ribonucleoprotein (RNP) complexes to specific target genes by interacting with them (Grote *et al*., 2013). Another example where these guide lncRNAs are important for the proper localization of the chromosome-modification complexes is in PRC2 and its subsequent repression of gene expression by the lncRNA fetal-lethal non-coding developmental regulatory RNA (*FENDRR*), which serves to bring PRC2 in close association to the promoters of genes linked with the build-up and differentiation of the lateral mesoderm lineage, such as forkhead box F1 (*FOXF1*) and paired-like homeodomain2 (*PITX2*) genes (Grote *et al*., 2013). Enhancer regions usually produce enhancer RNAs (eRNAs) and these eRNAs may work by modulating chromatin interactions also called 3-dimensional (3D) organization of DNA. Long non-coding RNAs may also work as “tethers” if they are not released from the enhancer regions when performing their role, thereby tethering the interacting proteins to enhancer regions (Li *et al*., 2014). Long non-coding RNAs can also encode short peptides, which may also play vital role in biological processes (Ingolia *et al*., 2011). As more research are being carried out to probe and unveil other functions of lncRNAs, there is no doubt, that in the near future, more functions of lncRNA will be reported in the fight against human diseases. Figure 2.2 below shows the mechanisms of lncRNAs.

### Cis and Trans-Acting lncRNA

Emerging evidence have shown that long non-coding RNAs can regulate gene expression by forming triplex structures either in cis-acting manner (on neighboring genes) or in trans-acting manner (on distant genes) (Guttman and Rinn, 2012; Guil and Esteller, 2012). Cis and trans-encoded RNAs are the most largely studied non-coding RNAs. While the first transcripts are in cis natural antisense relative to the targeted mRNA, the second transcripts in genomic regions are far from the targeted mRNA, therefore only cis-encoded RNAs have a perfect base pairing with the targeted mRNA (Azhikina *et al*., 2015). In the cis-acting manner, lncRNAs are usually transcribed from the promoter regions of target genes, form a triplex with promoter DNA sequences, and regulate gene expression while lncRNAs specifically target binding sites through triplex formation with distant DNA sequences in the trans-acting manner. While PARTICLE, Khps1, FENDRR, pRNA of rRNA gene, and pRNA of *DHFR* forms cis, MEG3 and HOTAIR forms trans conformations (Yue *et al*., 2016). Figure 1.3 indicates gene regulatory functions in cis or trans lncRNA.

**Fig 1.1:**
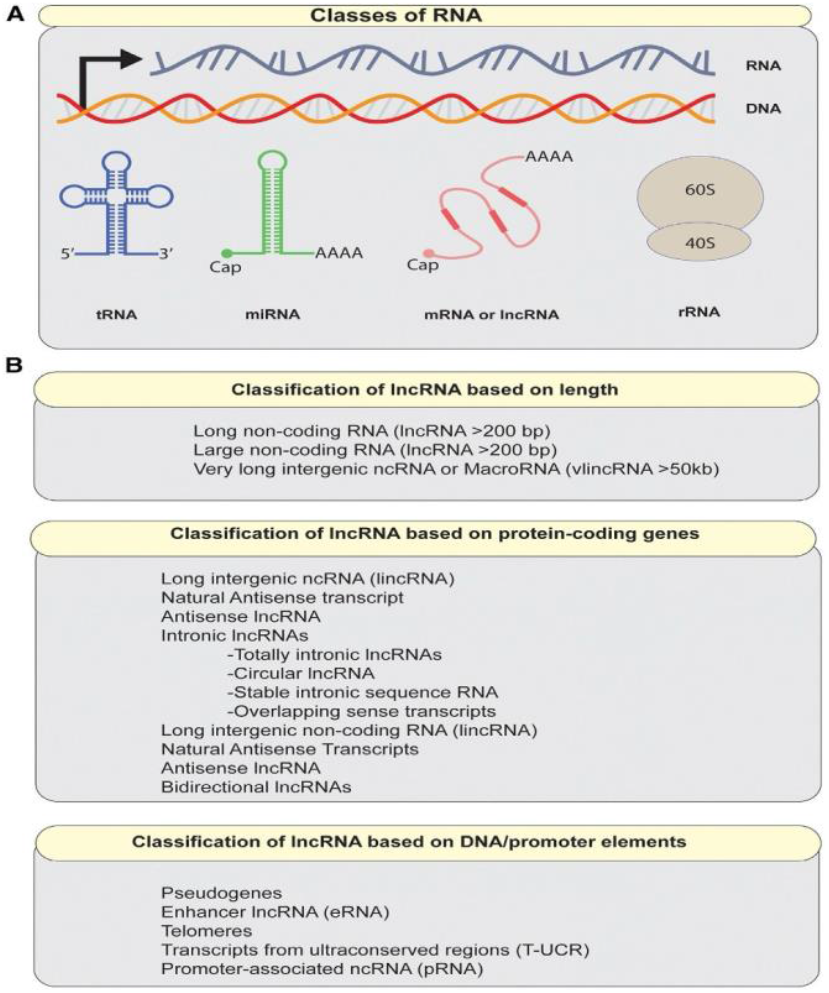
Types of RNA in mammalian cells. A; A main classes of RNA. Non coding RNAs may resemble mRNA structurally, but somecan have distinctive features, for example lack of polyA tail. B; Classes of noncoding RNAs >200 bp. qpcr stands for quantitative polymerase chain reaction (Sallam et al., 2018).

**Fig 1.2:**
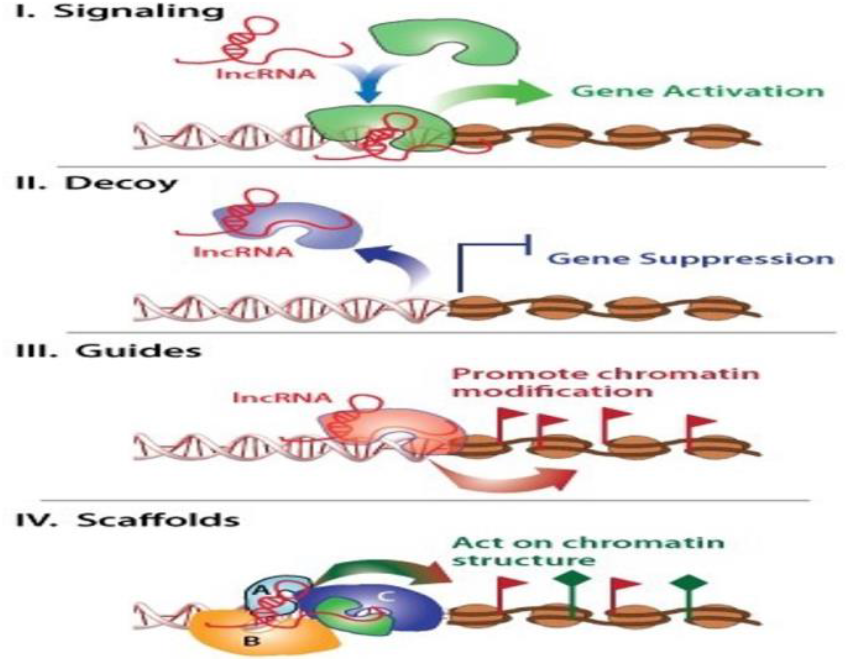
The molecular mechanisms of long non-coding RNAs. (Wang and Chang, 2011).

**Fig 1.3:**
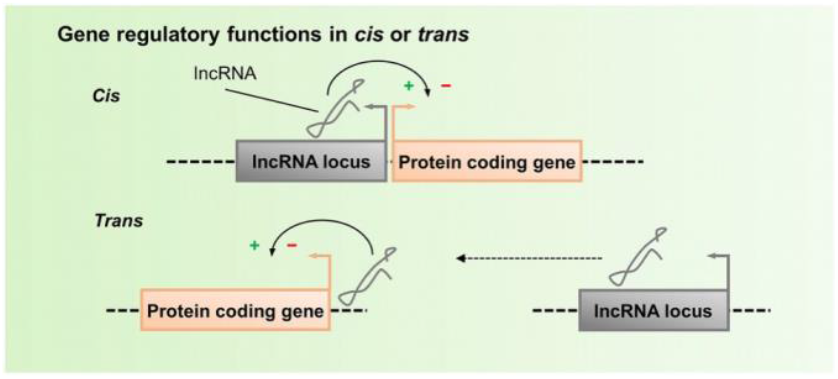
Gene regulatory functions in cis or trans lncRNA (Florian Kopp, 2019)

### RNA:DNA Triplex Forming Long Non-Coding RNA and their Structure

Several long non-coding RNAs (lncRNAs) have been verified experimentally by sequencing and other computational methods to form RNA:DNA triplex structures. These lncRNAs that have the potential of forming triplex structures include, HOTAIR, MEG3, PARTICLE, DACOR1, MIR100HG, FENDRR, ANRIL, TUG1, MALAT1, LINC00599, TINCR, NEAT1, roX2, DHFR, OTX2-AS1, Xist, SNHG16, ATXN8OS, BCYRN1, TERC, Khps1, among many others (Kazimierczyk *et al*., 2020). Recently, studies have shown that MEG3 lncRNA exerts their function by interacting with enhancers close to SMAD2, TGFBR1 and TGFB2 by forming triplex structures (Mondal *et al*., 2015). Mondal *et al* reported that by binding to distal regulatory elements, MEG3 is able to regulate the activity ofTGF-β genes. The GA-rich sequences of MEG3 binding sites can form RNA– DNA triplex structures with MEG3 which makes it easier to modulate chromatin activity.

MEG3 binding sites have GA-rich sequences (Mondal *et al*., 2015). HOTAIR lncRNA has been reported to interact with more than 900 genomic regions close to the binding sites of EZH2 and SUZ12 in human cancer cells (Chu *et al*., 2011). It has been reported HOTAIR lncRNA binds with DNA regions in a trans manner thereby recruiting PRC2 and LSD1-CoREST chromatin modifying complexes, resulting to transcription repression (Angrand *et al*., 2015). By forming RNA–DNA triplex structures, HOTAIR lncRNA is able to exert their function on the differentiation of mesenchymal stem cells (MSCs) and its association with senescence-associated DNA-methylation (DNAm) (Kalwa *et al*., 2016). O’Leary *et al* reported that the antisense lncRNA PARTICLE takes part in RNA-DNA triplex structure formation and is expressed and transcribed from the promoter of MAT2A when cells are exposed to low-dose of irradiation (O’Leary *et al*., 2017, O’Leary *et al*., 2015). By forming RNA:DNA triple helices, FENDRR lncRNA recruits PRC2 thereby interacting with Trithorax group and Mixed lineage leukemia (TrxG/Mll) complex (Grote and Herrmann, 2013). The lncRNA ANRIL forms RNA:DNA triple helices by binding with the DNA in a cis manner, thereby recruiting PRC1 and PRC2, leading to transcription repression (Angrand *et al*., 2015). For dosage compensation in mammalian females, the Xist lncRNA (X-inactivation specific transcripts) forms RNA:DNA triple helices with DNA regions and later recruits PRC2 to drop H3K27me3 repressive histone modification marks along the X-chromosome, thereby inactivating the marked copy (Zhao *et al*., 2008; Jeon and Lee, 2011). The formation of RNA:DNA triple helices between MIR100HG lncRNA and DNA regions plays an important role in triple-negative breast cancer (TNBC) progression by regulating p27 gene (Wang *et al*., 2018). The identification of Tug1-binding element (TBE) upstream of the *Ppargc1a* gene and its subsequent binding with Tug1 lncRNA have been linked to increase in *Ppargc1a* promoter activity (Jianyin *et al*., 2016).

Research has shown that a direct interaction between Tug1 and PGC-1α regulates mitochondrial bioenergetics in podocytes in the diabetic milieu (Jianyin *et al*., 2016). MALAT1 lncRNA carrying ENE and A-rich tract have also been reported to increase the levels of intronless β-globin reporter RNA (Jessica *et al*., 2012). An antisense RNA known as khps1 has been confirmed to form RNA:DNA–DNA triplex structures at the promoter region of Sphingosine kinase 1 (SPHK1), thereby regulating SPHK1 gene through histone acetylation that subsequently result in increase in cell proliferation activity (Postepska-Igielska *et al*., 2015). The role of NEAT1 lncRNA in leukemia and ovarian cancer through the formation of RNA:DNA triple helices have been revealed. It does this by regulating ADARB2 expression through protein sequestration into paraspeckles and also plays an important role in the knockdown results in inhibition of cell growth (Ke *et al*., 2016; Hirose *et al*., 2014). For a given RNA sequence, *R = r*_*1*_*r*_*2*_*···r*_*n*_, and DNA sequence *D = d*_*1*_*d*_*2*_*···d*_*0*_, the triplex detection problem is the search for (maximum length) substrings *r*_*i*_…*r*_*j*_ *and d*_*u*_..*d*_*v*_ from *r* and *d* with minimum length *l* (). Figure 1.4 below shows long non-coding RNAs in prostate cancer.

**Fig 1.4:**
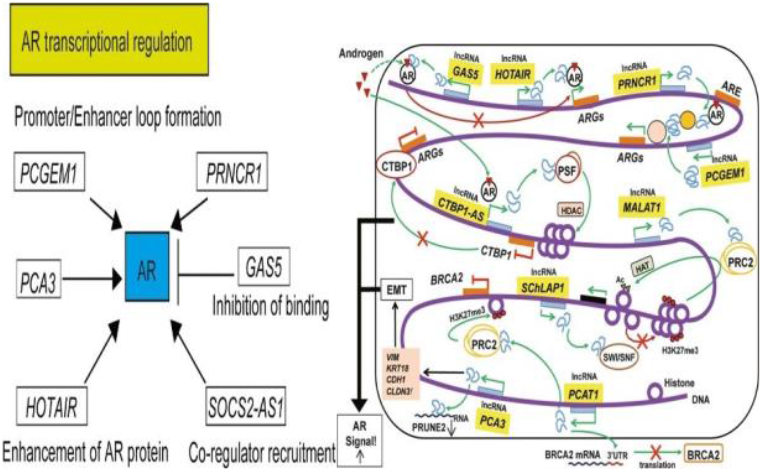
Long non-coding RNAs in prostate cancer (Misawa *et al*., 2017).

**Fig 1.5:**
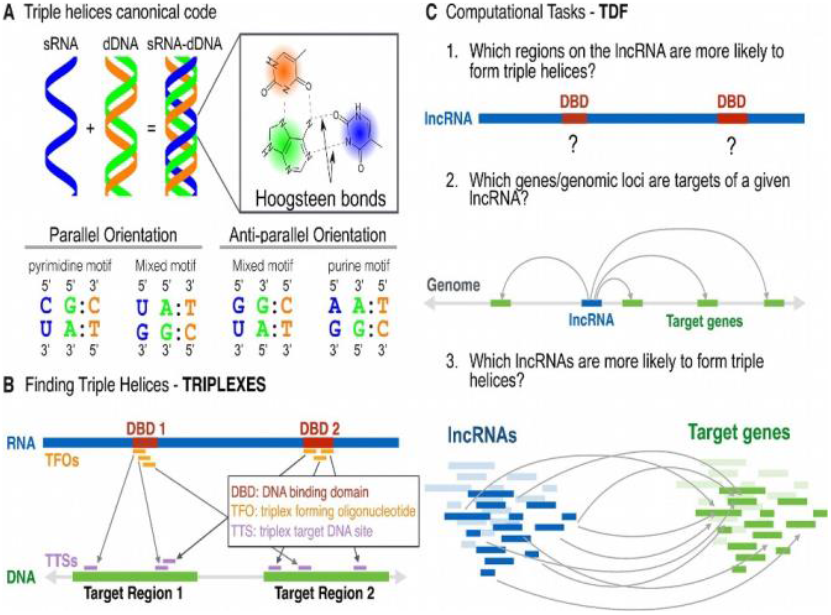
The computational framework of *triplexes* and TDF. (A) Triplexes are formed by binding of single-strandedRNA (blue) with a purine-rich strand (green) of a double-stranded DNA through Hoogsteen base pairing rule. For the formation of a triplex in the parallel orientation, a pyrimidine or mixed motifs are needed, but the anti-parallel orientation needs a purine or mixed motifs. (B) For a given RNA and DNA sequence, T *triplexes* identifies candidate triple helices with a minimum size and maximum number of mismatches following one of the canonical codes. Eachtriplex is formed by one RNA sequence (triplex forming oligo – TFO) and a DNA region (triple target sites – TTS). Contiguous regions with overlapping TFOs (marked in red) define a DNA-binding domain. (C) TDF performs statistical tests by combing predictions from *triplexes* to answer the following questions: (1) which regions of an RNA (DBD) are more likely to form triple helices with particular DNA target regions? (2) Which DNA regions (target genes) are more likely to be targeted by the RNA? and (3) which lncRNAs are more likely to form triple helices in a set of target DNA regions? (Kuo *et al*., 2019).

### Long Non-Coding RNA (lncRNA) Interactome

Long non-coding RNAs are capable of interacting with DNA, messsenger RNAs (mRNA), proteins, peptides and micro RNAs (mRNA) – the so called competitive, endogenous RNA (ceRNA) hypothesis.

### Interactions of lncRNA with miRNAs - the competitive, endogenous RNA (ceRNA) hypothesis

Vast novel evidence indicates that miRNAs interacts with lncRNAs by binding to specific mRNA 3’ UTR regions thereby modulating the expression of these genes (Krol et al., 2010). Large amounts of experimental data suggests that ncRNAs interacts with miRNAs and this evidence was named as competitive, endogenous RNA (ceRNA) hypothesis (Smillie *et al*., 2018; Yay *et al*., 2014; Salmena *et al*., 2011).

### Pairing LncRNAs with Messenger RNAs (mRNA)

Bioinformatics tools such as LncTar and RNAplexn have given us knowledge on the kind of interaction between lncRNA and mRNA (Li *et al*., 2015; Tafer and Hofacker, 2008). LncRNA interacts with mRNA and the pre-mRNA by direct base pairing, which then plays a vital role in modulating mRNA splicing and editing may participate in regulating translation by affecting mRNA splicing and editing together with its stability (Romero-Barrios *et al*., 2018; Karapetyan *et al*., 2013). Long non-coding RNA can regulate pre-mRNA splicing by either blocking spliceosome assembly involving intron-exon junction or by becoming the target for splicing factors (Romero-Barrios *et al*., 2018). More experimental evidence is needed to confirm lncRNA– mRNA interaction and its role in alternative splicing as research proposed the involvement of MALAT1 lncRNA in modulating alternative splicing of pre-mRNA (Tripathi *et al*., 2010).

### LncRNA*–*DNA Interactions

There are a lot of approaches depicting the interaction of lncRNAs and DNA regions (Quinodoz and Guttman, 2014). It has been demonstrated that LncRNAs can form RNA-DNA-DNA triple helix (Li *et al*., 2016; Shaw and Arya, 2008). For example, research has demonstrated the regulatory role of DHFR lncRNA in suppressing the transcription of Dfhr mRNA through the formation of triple helices with the DHFR promoter, thus resulting in the binding of DHFR lncRNA to the TFIIB and blocking the formation of transcription initiation complex (Deng *et al*., 2009; Blume *et al*., 2003).

### LncRNA*–*Protein Interactome

Novel evidence has emerged that long non-coding RNAs (lncRNAs) weight ranging from hundreds to tens of thousands of nucleotides plays vital roles as either signals, decoys, guides, and scaffolds for majority of proteins (Sun *et al*., 2017; Wang and Chang 2011).

Today, there are three approaches to study the interaction between lncRNAs and proteins. These include the immunoprecipitation method, crosslinking and immunoprecipitation (CLIP) method and Next Generation Sequencing (NGS) approach. The immunoprecipitation approach has been applied in the preparation of RNA-protein complexes while on the other hand, crosslinking and immunoprecipitation method (for example HITS-CLIP, PAR-CLIP, and iCLIP) works to map out proteins and protein binding sites interacting with a sequence of lncRNA (Jalali *et al*., 2018; Garzia et al., 2017; Yoon and Gorospe, 2016). To accelerate the the analysis of RNA-protein complexes, Next Generation Sequencing (NGS) have been rapidly applied in generating data and studying the interaction between lncRNAs and protein. Computational approaches that seek to predict the structure and function of lncRNA-protein by screening thousands of publicly available data such as annotated sequences and structural motifs (for example, RNA-binding sites) are gaining grounds in recent years.

### LncRNA-Peptides and small weight compounds Interactome

Research has shown that small portions of lncRNAs in the cytosol can code for small peptides whose amino acid composition is less than 100 (Chugunova *et al*., 2018; Matsumoto and Nakayama, 2018; Zhu *et al*., 2018). For example, a 16.5 kb long skeletal and muscle-specific lncRNA known as LINC00948 has been confirmed to encode for myoregulin (a 46 amino acid peptide), thus modulating the biosynthesis of skeletal muscles by interacting with sarcoplasmic reticulum Ca 2+-ATPase (SERCA) (Anderson *et al*., 2015), lysosomal v-ATPase (Matsumoto *et al*., 2017) or removing SERCA inhibitors (Nelson *et al*., 2016). Thetask of profiling lncRNA-origin peptides is a difficult one since *in silico* methods predicts large numbers of Open Reading Frames (ORF), meanwhile very little of them are encoded to yieldproteins. Ruiz-Orera *et al* estimated that only about out of the transcribed lncRNAs, only about 23% of them translated to proteins (Ruiz-Orera *et al*., 2016). The use of bioinformatic algorithms and experimental confirmation has helped in profiling peptide expression. Large amounts of RNAs such as rRNA, mRNA, ribozymes, riboswitches, and aptamers forms small structures containing “pockets” that are capable of binding with compounds with small molecular weights (Warner *et al*., 2018; Gestelanf *et al*., 2005). Figure 1.6 shows the interaction of human lncRNA with cellular biomolecules.

**Fig 1.6:**
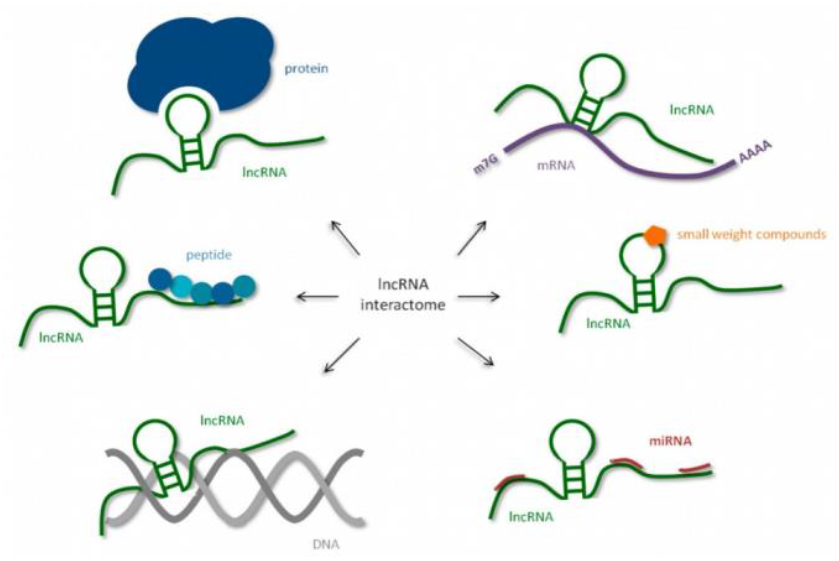
Human long non-coding RNAs (lncRNA) interactome, interaction of human lncRNA with cellular biomolecules (Kazimierczyk *et al*., 2020).

### Sequencing Technologies for Identifying Long Non-Coding RNA

Either examining mammalian cells, tissues, or serum samples, genome-wide transcriptomic methods is the basis of profiling long non-coding RNAs. Different computational techniques such as classical machine learning and deep learning methods can also identify long non-coding RNAs from RNA-seq data. Generally, RNA-seq data can be analyzed through two different ways: de novo assembly methods by using software packages such as Bridger (Chang *et al*., 2015), SOAP2 (Li *et al*., 2009), Oases (Schulz *et al*., 2012), or Trinity (Grabherr *et al*, 2011), or by alignment against a known standard of defined lncRNAs. The alignment against a known standard of defined lncRNAs method is rapidly used today due to increasing lncRNA annotations.

Even though the understanding about histone modification marks that defines active transcription can be helpful, it is not needed in screening lncRNAs by sequencing approaches. At its promoter, H3K4me3 marks a subunit of lncRNAs showing an element of mammalian conservation while H3K36me3 does so along the region where it is transcribed (Kaikkonen *et al*., 2011; Guttman *et al*., 2009). This chromatin signature is sometimes known as K4k36. Even though histone modification marks can help in identifying lncRNAs, more research is being carried out to know things that differentiate long non-coding RNAs from other non-coding RNAs that regulates transcription for example, enhancer RNAs. The main work of chromatin-binding patterns to define transcription regulators has limitations even though it is one among the core factors that differentiate lncRNAs from other transcription regulatory elements (Li *et al*., 2016; Li *et al*., 2013).

Despite RNA-seq being the most widely used sequencing technique for RNA screening, there are other methods that can be used in detecting RNAs and these include;

i. serial analysis of gene expression (SAGE) and paired-end tag–expression approaches thattargets polyA tail (Philippe *et al*., 2014; Fullwood *et al*., 2009);
ii. GrRO-seq: is a nuclear run-on assay coupled to deep sequencing to estimate nascent transcription (Gardini, 2017; St Laurent *et al*., 2015).
iii. direct RNA sequencing, that requires sequencing of undegraded RNA bypassing library preparation (Ozsolak *et al*., 2009); and
iv. cap analysis of gene expression (CAGE), which profiles RNAs with 5′ cap and can accurately map the 5′ transcript end which makes it more advantageous (Takahashi *et al*., 2012).

Today, several screening microarrays are available in commercial quantities that can help in profiling protein-coding and non-coding transcripts from human or mouse samples (Zhu *et al*.,2017; Shi and Shang, 2016; Chen *et al*., 2015).

Microarrays have proven to be fast and efficient which may be advantageous over RNA-seq.

Nuclear lncRNAs may determine transcriptional products in different ways, which include gene expression regulation by interacting with transcription factors, histone modulation, and its regulatory role in the splicing process and subsequent transport of mRNA out of the nucleus (Wang KC and Chang HY, 2011).

Recently, different sequencing approaches have been developed that can assist in profiling lncRNA interactions (Figure 2.7; 2.7.1 & 2.7.2). The first chromatin affinity approach (chromatin isolation by RNA purification sequencing - ChIRP) to characterize lncRNA interactions was developed by Chu *et al* (Chu *et al*., 2012; Chu *et al*., 2011). Several other research later validated this technique (Sallam *et al*., 2016). Chromatin isolation by RNA purification sequencing (ChIRP) method normally captures lncRNA of interest and probes its interaction by tiling complimentary antisense oligos. By so doing, ChIRP-Seq technique is able to define specific DNA binding sites of a particular lncRNA and by coupling this with mass spectroscopy, it helps in characterizing interacting protein partners. By this approach, we have been able to understand the complex interactions of Xist lncRNA and its relationship with hnRNP K and Spen (Spen family transcriptional repressor), which are needed for mRNA processing (Chu *et al*., 2015). ChIRP-Seq has been applied in studying the interactions between RNA– RNA (Chu *et al*., 2012). RNA antisense purification - RAP (Engreitz *et al*., 2015) and capture hybridization analysis of RNA targets - CHART (Simon *et al*., 2011) are also used in studying the binding motifs based on biotin-tagged oligonucleotide retrieval. While RAP uses longer probes (≈90 base pair designed with 5′ biotin tags), ChIRP uses 20 nucleotide probes with 3′ tags. CHART-Seq also uses 20 mer olignonucleotides, although what it does is to first test individual olignonucleotides that targets ncRNA regions that are accessible by using RNase sensitivity. ChOP-Seq is another sequencing technique that has been applied in studying the kind of interaction lncRNA has with other biological molecules. ChOP-Seq has been applied in studying the complex interactions of kcnq1ot1 lncRNA that take part in the regulation of genes within the *Kcnq1* imprinting domain (Zhang *et al*., 2014).

**Fig 1.7:**
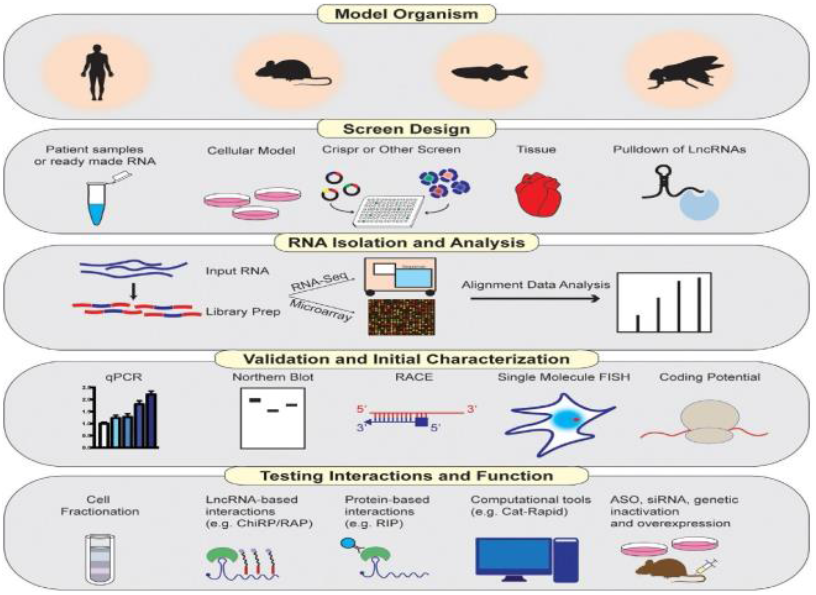
Pipeline for long non-coding RNA (lncRNA) discovery and characterization (Sallam *et al*., 2018).

**Fig 1.7.1:**
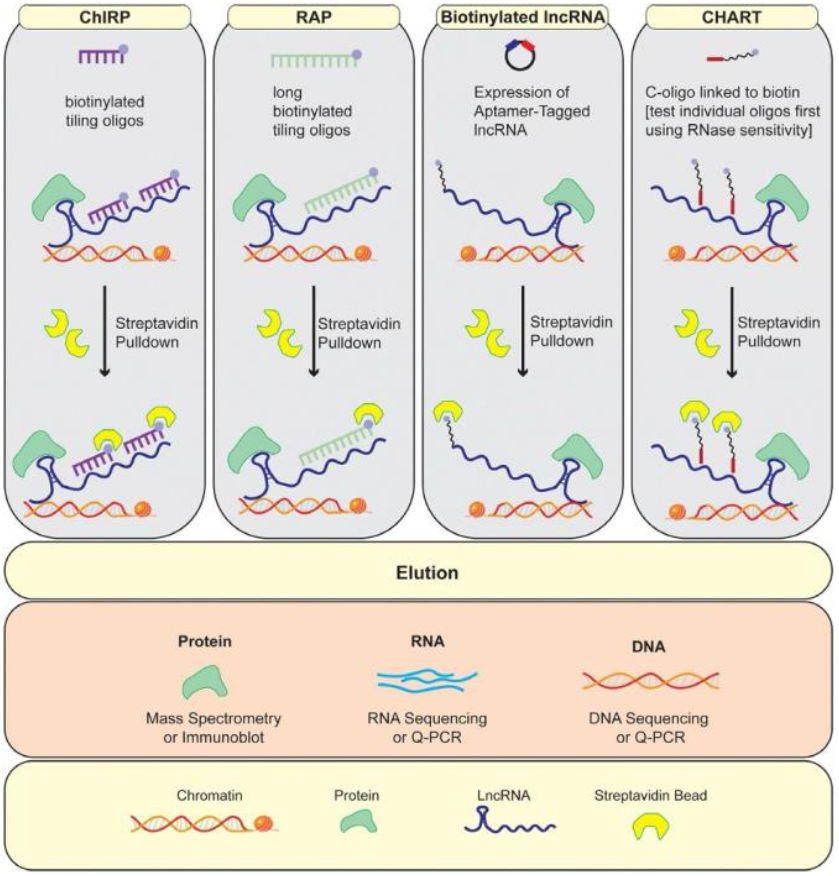
Summary of techniques used to probe a long non-coding RNAs (lncRNA) interactome (Sallam*et al*., 2018).

**Fig 1.7.2:**
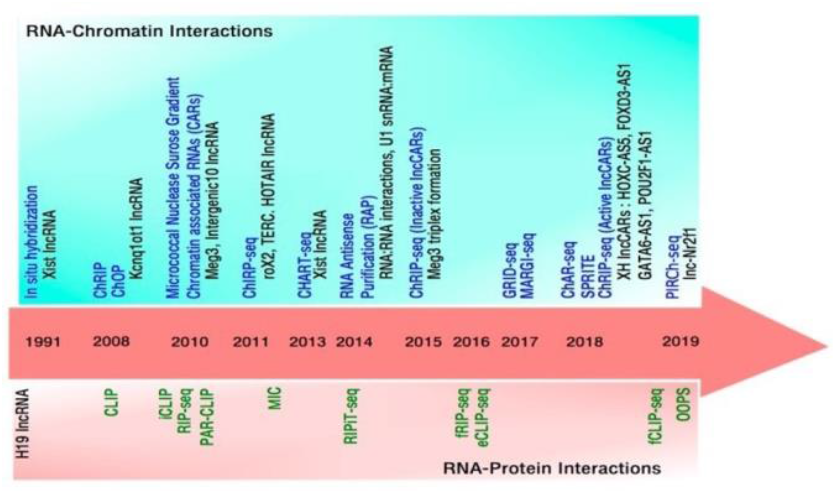
Timeline of technological advances to study RNA-Protein and RNA-Chromatin interactions. Upper panel (light sea-green box) shows in chronological order the prominent methods (in blue) to detect RNA-chromatin interactions. Examples of some of the functionally verified lncRNAs from each of these studies are shown (in black) below the corresponding method. Middle panel shows the year in which these methodologies were reported Lower panel (light pink box) likewise shows in chronological order methods (in green) to recognize RNA-proteins interactions (Mishra and Kanduri, 2019).

### Base Pairing Rules-Related Mathematics Statistics for Predicting DNA:RNA Triplex Structures

Computational approaches play an important role in recognizing lncRNA triplex structures. Earlier computational approaches did not mark out triplex forming RNA regions, because they were based on searching for purine rich DNA regions (Gaddis *et al*., 2006; Goñi *et al*., 2004). Subsequently, another algorithm known as Triplexator was developed which efficiently recognized lncRNA triplex structures in large DNA sequences (Buske *et al*., 2012). Triplexator was able to categorize all regions of RNA and DNA sequences that would take part in the formation of triplex structures with size larger than l basepair and with k maximum mismatches. Today, Triplexator is a widely used computational tool that recognizes tens of thousands of triplex structures for a single lncRNA, although it provides little statistics to identify significant triple helices. Triplexator is able to recognize vast amounts of lncRNA triplex structures including; HOTAIR, MEG3, PARTICLE, DACOR1, MIR100HG, FENDRR, ANRIL, TUG1, MALAT1, LINC00599, TINCR, NEAT1, roX2, DHFR, OTX2-AS1, Xist, SNHG16, ATXN8OS, BCYRN1, TERC, Khps1, among others (Kazimierczyk *et al*., 2020). Other computational approaches for recognizing triplex structures include, LongTarget, Triplex-Inspector and Triplex Domain Finder (He *et al*., 2014 Buske *et al*., 2013). LongTarget is a web-based technique which estimates only a single DNA region at a time and has been shown recently to be significantly slower than Triplexator and as a result it hinders the analysis of large amounts of RNA or DNA regions (Antonov *et al*., 2018). Triplex-Inspector is another computational method for evaluating triple helices. It is for sequence-specific ligand design and suitable target selectionwhich increases the accuracy of triplex-based genomic manipulations (Buske *et al*., 2013). The algorithm takes a genomic locus of interest and searches for all putative triplex target sites, leveraging our exhaustive search algorithm Triplexator (Buske *et al*., 2012). The last computational approach we mentioned above is the Triplex Domain Finder (TDF) which recognizes the triplex-forming potential between lncRNA and DNA regions such as promoters of differentially regulated genes after knockdown of the lncRNA (Kuo *et al*., 2019).

### Classical Machine Learning Methods in Predicting DNA:RNA Triplex Structures

Recently, vast amounts of biological data have posed a huge challenge in understanding and interpreting them correctly. Having knowledge of these big data will not only answer questions about the etiology of animal and plant diseases; it provides an opportunity to appreciate technological advancements in computational sciences. Data generating technologies such as next generation sequencing (NGS) have enabled us to screen large amounts of omics data such as genomic data, gene expression and small RNA abundance data, epigenetic modification data, protein binding motifs, and chromosome conformation in a as shown in (Figure 2.). These omics data come with its own challenges such as heterogeneity, noisy and highly dimensional (Zitnik *et al*., 2019; Altman and Krzywinski, 2018). Today, machine learning/deep learning has become a powerful computational tool in the field of biology. Machine learning-deep learning learns new features from big omics data, thus answering penitent research questions. Classical machine learning tools such as support vector machines, random forests, decision trees, etc have been applied variously on omics data in the past three decades. While most classical machine learning methods are supervised learning, deep learning on the other hand is unsupervised learning. These methods have proven over the years to be fast, efficient and easy to understand and interpret. Figure 1.8 shows machine learning workflow for analyzing complex biological data.

**Fig 1.8:**
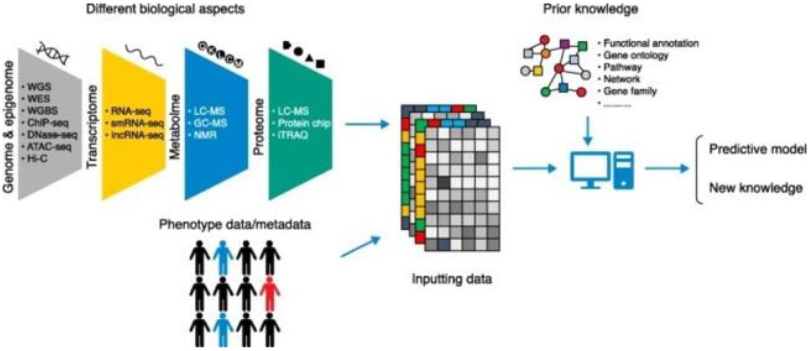
Machine learning workflow for analyzing complex biological data. High-throughput data generation techniques for different biological aspects are shown (*left*). *ATAC seq* assay for transposase-accessible chromatin using sequencing, *ChIP-seq* chromatin immunoprecipitation sequencing, *Dnase-seq* DNase I hypersensitive sites sequencing, *GC-MS* gas chromatography-mass spectrometry, *LC-MS* liquid chromatography–mass spectrometry, *lncRNA-seq* long non-coding RNA sequencing, *NMR* nuclear magnetic resonance, *RNA-seq* RNA sequencing, *smRNA-seq* small RNA sequencing, *WES* whole exome sequencing, *WGBS* whole-genome bisulfite sequencing, *WGS* whole genome sequencing, *Hi-C* chromatin conformation capture combined with deep sequencing, *iTRAQ* isobaric tags for relative and absolute quantification (Xu and Jackson, 2019).

### Support Vector Machines (SVM)

The support vector machine (SVM) is a computational algorithm that seeks to learn from the data via pattern recognition. This computational tool has been applied on several big omics data including DNA sequences, protein sequences, microarray expression profiles, protein-protein interaction networks, and tandem mass spectra among others (Zitnik *et al*., 2019).

Binary classification is the easiest way of making predictions, thus discriminating between objects that belong to one of two categories, either positive (+1) or negative (−1). SVMs has two main components of solving this task, either by large margin separation or by kernel functions (Ben-Hur *et al*., 2008). Classification of points in two dimensions forms the basis of large margin separation and is done by drawing a straight line and naming points on one side as positive (+) and on the other side as negative (−) (Ben-Hur *et al*., 2008). This can be expressed mathematically as:

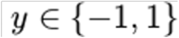

where y is the output while 1 is the positive class and −1 is the negative class. Kernel functions usually take two vectors of any dimension as its input and gives an output score that shows similarities between input vectors (Ben-Hur *et al*., 2008). This means that if the dot product is large, then it means that the vectors are similar, but if the dot product is small, it therefore means that the vectors are different. Mathematically, a Kernel function can be written as:

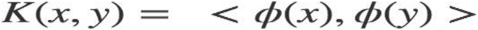

where **x** and **y** are input vectors, ***ϕ*** is a transformation function and <, > denotes dot product operation. ***ϕ*** only maps the input vector to itself in dot product function.

### Random Forests

Random forests (RF) is a powerful algorithm that can be applied on both small and big data. Since it can study interactions between features and interpret its correlation, it is applied on high-dimensional omics data (Wouter *et al*., 2013). Random forest models are fast to train, non-parametric, difficult to over-train, very robust to outliers and noise and can be trained without hyperparameter tuning, even though tuning hyperparameters is good for training machine learning-deep learning models (Wouter *et al*., 2013; Breiman, 2001). Random forests can train an ensemble of decision trees singly according to their samples, variables and class designation (Wouter *et al*., 2013). The name random forest is derived from the idea of using random subset of variables and samples to build every tree. Mathematically, for each classifier; *hk(****x***) ≡ *h(****x***|*ΘΘk)* is a predictor of *n*

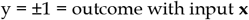

### Decision trees (DTs)

Decision trees (**DTs**) are supervised learning and they are fast to train, non-parametric, simple to understand and interpret, can work on small amount of data, and have been applied on regression and classification tasks in biology (Carl and Steven, 2009). The aim is to generate a model that can predict the value of a target variable. It does this by learning simple decision rules gathered from the data features (Carl and Steven, 2009). The application of decision trees in biology spansfrom predicting splice sites to assigning functions to different protein (Carl and Steven, 2009).

### Deep Learning Methods in Predicting DNA:RNA Triplex Structures

In recent years, deep learning (DL) architectures have recorded successes in speech recognition, image recognition and language processing (LeCun Y *et al*., 2015). Deep learning is a subset of machine learning algorithm. Today, different deep neural networks like convolutional neural networks and recurrent neural networks have been applied on different biological tasks including splice site detection, protein-ligand scoring, protein-protein interaction prediction, protein secondary structure prediction, protein contact map prediction, compound toxicity, liver injury prediction, RNA-DNA triplex helix structure recognition, protein design drug repurposing among others (Yongqing *et al*., 2020). Since they can learn new important features from the ever increasing heterogeneous and complex omicsdata via interactions within the layers, it has helped in the diagnosis, treatment, and classification of different complex diseases. Deep neural network’s goal is to calculate a mathematical function say *f* that takes a number of inputs called, *x*, to their respective outputs known as, *y*, such that *y* = *f*(*x*). For a simple feed-forward network, *y* = *f*(*x;w*) = *LN*(*LN*-1(….*L*1(*x*)) can be defined as a composition of N nonlinear transformations *Li*(1<=*i*<=*N*) where each function *Li* corresponds to a hidden layer activation, while *w* is the learnable weight contained in all filter bank layers that are updated in the course of the training process (Martorell-Marugán *et al*., 2019).

### Convolutional Neural Network (CNN)

Convolutional neural networks (CNNs) have become a conventional tool for the analysis of omics data as feedforward networks (Waseem *et al*., 2017). The performance of CNNs largely depends on the technical know-how on tuning both internal and external parameters. CNNs have worked well on DNA sequences over the years, hence providing answers to numerous biological questions and paving way for disease sub-typing and treatment. Convolutional layer, pooling layer (subsampling), and fully connected layer are the building block layers of a convolutional neural network architecture despite coming in different forms (Waseem *et al*., 2017). The convolutional layers usually extract features and learns how the neurons in this layer are represented as feature maps (Waseem *et al*., 2017).

### Recurrent Neural Network (RNN)

Recurrent neural networks (RNNs) basically make use of sequential data. The network works such that the decision of the first output say time t1 affects the decision of the second output say time t2. RNN have two input sources, the present source and the preceding source, and the coming together of these two input sources determines the kind of respond to a new data (Martorell-Marugán *et al*., 2019). Evidence from research have shown that recurrent neural networks (RNN) performs better on time-series data than other machine learning and deep learning architectures (Shen *et al*., 2018).

### Residual Neural Network (ResNN)

Residual neural networks (ResNN), was first reported by He *et al* in 2016 (He *et al*., 2016) and since then, it has outperformed other machine learning and deep learning architectures. The performance of ResNN in practice can be credited to the skip connections between layers that helps in the propagation of the gradient round the network. Aside from solving the gradient problem in deep networks, the skip connections establishes a dependency between variables in different layers that can be seen as a system state (Ciccone *et al*., 2018; Chang *et al*., 2018; Haber and Ruthotto, 2017; Lu *et al*., 2017; Liao and Poggio, 2016; Ruthotto and Haber, 2018; Chaudhari *et al*., 2017).

### Multilayer Perceptron (MLP)

A multilayer perceptron (MLP) is also a class of feedforward artificial neural network and it comprise of at least three layers of nodes namely; an input layer, a hidden layer and an output layer. Apart from the input nodes, each node is a neuron that makes use of a nonlinear activation function. For model training, multilayer perceptron (MLP) uses a supervised learning method known as backpropagation. The distinguishing factor between MLP and a linear perceptron is its multiple layers and non-linear activation. This means that it can distinguish a data that is not linearly separable. Multilayer perceptron (MLP) has been widely applied on omics data. For example, multilayer perceptron (MLP) has been utilized for the analysis of immunohistochemical scores of tissue microarrays for breast cancer biomarker discovery (McKenna *et al*., 2013).

### Long-/Short-Term Memory (LSTM)

The main problem of RNNs is that of vanishing gradients, thereby significantly limiting its ability to work with long sequences (Martorell-Marugán *et al*., 2019). To solve this problem, Long-/short-term memory (LSTM) and gated recurrent units partially address this issue (Martorell-Marugán *et al*., 2019). Long-/short-term memory (LSTM) seeks to maintain the error that can be back-propagated via time and layers after learning over several steps.

### Methods of Hyperparameter Optimization and Others

Hyperparameters are basic building blocks that occurs in all machine learning models to check their structure and performance. Some of the external hyperparameters that are optimized for better accuracy in model training include; learning rate, grid search, training epouchs, Adams optimization algorithm, etc. The learning rate, also known as *step size* defines how much we need to configure the model based on the estimated error when weights are calculated and this is usually done in the range of 0.0 to 1.0 (Park *et al*., 2020). Tensorflow and Pytorch works best at a learning rate of 0.001 while in Natural Language Processing, 0.002 and 0.003 has been reported to perform best. Grid search is an automatic hyperparameter optimization technique that is user-friendly, but may not work efficiently when the number of parameters is large. Epochs specifies the number of rounds of the entire training dataset the machine learning algorithm and deep neural network has completed. Using many epochs to train the model may lead to overfitting, thus making the model to learn even the noise from the parameters. Adam is an optimization algorithm used in place of stochastic gradient descent to update model weights in the training data (Diederik and Jimmy, 2015).

### Activation Function

Activation functions are vital in constructing neural networks. Today, ReLU activation function is the default activation function for hidden layers while the output activation function depends on the task you are predicting. Rectified Linear Activation (**ReLU**), Logistic (**Sigmoid**), and Hyperbolic Tangent (**Tanh**) are widely used as hidden layer activation function for deep learning models. Recurrent neural networks uses Sigmoid and Tanh activation functions. Most times, LSTM uses Sigmoid activation function for recurrent connections and Tanh activation as an output activation function. Below are the hidden layer activation functions widely used in MLP, CNN and RNN;

*Multilayer Perceptron (MLP)*: ReLU activation function.

*Convolutional Neural Network (CNN)*: ReLU activation function.

*Recurrent Neural Network (RNN)*: Tanh and/or Sigmoid activation function.

For the output layer, there are three commonly used activation functions, namely; Linear activation function, Logistic (Sigmoid) activation function, and Softmax activation function. Below is how output activation functions can be chosen in different tasks.

*Binary Classification:* One node, Sigmoid activation.

*Multiclass Classification:* One node per class, Softmax activation.

*Multilabel Classification:* One node per class, Sigmoid activation.

*Regression:* Linear activation function

### Performance Metrices

An important step for machine learning methods is the choice of metrics that can measure the real performance of models. Every performance metric has a particular feature and calculates properties that may be different from the results that has been predicted (Orozco-Arias *et al*., 2020). The performance of regression models can be determined by the following performance metrices; Mean Absolute Error (MAE), Mean Squared Error (MSE), Root Mean Squared Error (RMSE), and R^2^ (R-Squared) while the performance of classification models can be calculated by the following metrices; Accuracy, Confusion Matrix (not a metric but fundamental to others), Precision and Recall, F1-score, AU-ROC, and MCC (Orozco-Arias *et al*., 2020).

### Loss Functions

The loss function to use on any machine learning-deep learning tasks depends on the kind of task you want to solve. Different loss functions exists for different tasks as follow;

#### Regression Task

Here a real-value quantity is predicted where the output layer configuration is a one node with a linear activation unit and the loss function is mean squared error (MSE) (Wang *et al*., 2020).

#### Binary Classification Task

Here you usually classify an example as belonging to one of two the classes where the output layer configuration is a one node with a sigmoid activation unit and the loss function is cross-entropy, also known as Logarithmic loss. The task is designed as predicting the likelihood of an example belonging to class one (Wang *et al*., 2020).

### Multi-Class Classification Task

Here you classify an example as belonging to one of more than two classes where the output layer configuration is a one node for each class using the softmax activation function and the loss function is cross-entropy, also known as Logarithmic loss. The task is designed as predicting the likelihood of an example belonging to each class (Wang *et al*., 2020).

## Materials and methods

### Data Collection and Preprocessing

We installed biopython after which we cloned the GitHub repository of Zhang *et al*., 2020 to access the data to a hosted runtime provided by Google (Google Colab).

### Data Collection & Preprocessing for our model: Triplex Forming Prediction Model (TriplexFPM-HMM) for Histone Modification Marks-Based Feature

We took *2547* triplex positive DNA regions from Sentürk’s paper (GEO Accession No: GSM3417036), *3014* triplex negative DNA regions from Sentürk’s paper (GEO Accession No: GSM3417036) as our first negative data and *36597* promoter regions of DNA from Ensembl Release 104 of May 2021 as our second negative data.

### Triplex DNA Sites Prediction Positive Dataset for TriplexFPM-HMM

We took *2547* triplex positive DNA regions as a bed file from Sentürk’s paper (GEO Accession No: GSM3417036), we then removed all other columns apart from the chromosome number, start and end regions to enable us add histone modification marks.

### Triplex DNA Sites Prediction First Negative Dataset for TriplexFPM-HMM

In the same way, we also took *3014* triplex negative DNA regions as a bed file from Sentürk’s paper (GEO Accession No: GSM3417036) as our first negative data. Using pandas, we deleted all other columns and the header apart from the chromosome number, start and end regions to enable us add histone modification marks.

We took our 12 histone modification read counts from ENCODE database No. E123 cell line out of k562 cell lines (Inoue F. *et al*., 2017; Kazachenka *et al*., 2018), we converted the ‘tagalign.gz’format to ‘bam’ by following the instruction given on DeepChrome (Ritambhara *et al*., 2016). Wethen added our 12 histone modification marks to both our positive and negative data using bedtools multicov command.

Overall, we had *5561 DNA regions* and 12 histone modification features (H3K36me3, H3K27ac, H3K9me3, H3K4me3, H3K4me2, H3K9ac, H3K9me1, H3K4me1, H2AFZ, H3K27me3, H3K79me2, H4K20me1). *2547* DNA regions were Positive while *3014* were Negative DNA regions. After filtering out sequences greater than 2000 Bps, we decreased the number of regions from *5561 to 5473*.

**Fig 2.1:**
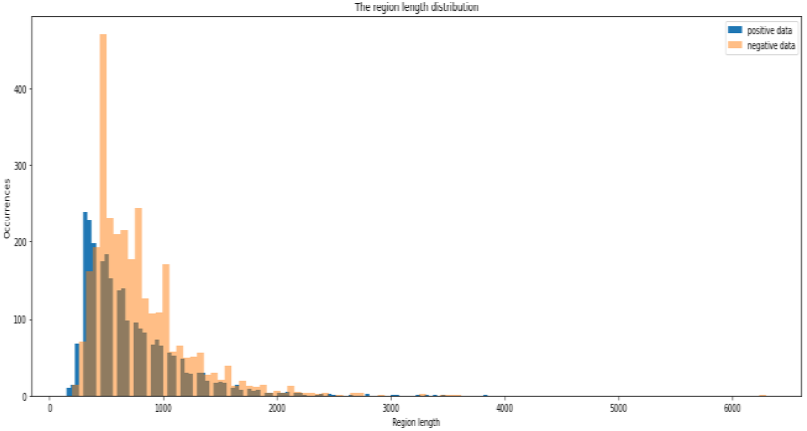
Triplex DNA site length distributions for positive & first negative dataset for TriplexFPM-HMM

### Model Construction for First Negative Dataset TriplexFPM-HMM

We splitted our filtered data *(5473)* into three sets; train *(0*.*8), validation (0*.*1) and test (0*.*1) sets* and trained MLP and CNN.

### Hyperparameter Tuning for First Negative Dataset TriplexFPM-HMM

We set other internal and external hyperparameters as follow: epochs (50), learning rate at (1e-2), input shape (12, 1) Kernel size (32), verbose (1) and used Keras-Tensorflow to train our models. We used six different metrices in assessing the performance of our models, these include: binary accuracy, precision, recall and AUC, F1-score, MCC. We also used ReLU activation as our hidden layer activation function and Sigmoid activation function as our output layer activation function for both MLP and CNN. We used Adam optimization algorithm as our optimizer and since it is a binary classification task, we also used binary cross entropy. We employed early stopping criteria on the optimization procedure when the values of the loss function on the validation set stopped decreasing.

### Triplex DNA Sites Prediction Second Negative (Promoters) Dataset for TriplexFPM-HMM

The NN training does not go well - the loss does not decrease through the training epochs. One of the reasons to this problem may be the faulty negative data that we used from the original paper. To handle this, we decided to adapt the Zhang *et al*., 2020 approach, where they used random promoter regions as their negative data.

We took *36 597* promoter regions of DNA from *Ensembl Release 104 of May 2021* as our second negative data (Kevin, *et al*., 2021). Our promoter regions are of Ensembl Regulation 104 of the human regulatory features (GRCh38.p13). We dowloaded only three columns; chromosome number, start and end regions from Ensembl after which we used pandas to remove the header and toadd *chr* to every chromosome number and saved it as a bed file.

We took our *12* histone modification read counts from ENCODE database No. E123 cell line out of k562 cell lines (Inoue *et al*., 2017; Kazachenka *et al*., 2018), we converted the ‘tagalign.gz’format to ‘bam’ by following the instruction given on DeepChrome (Ritambhara *et al*., 2016. We then added our 12 histone modification marks to our second negative data using bedtools multicov command.

Overall, we had *36 597 DNA promoter regions as our negative data* and *12 histone modification features* (H3K36me3, H3K27ac, H3K9me3, H3K4me3, H3K4me2, H3K9ac, H3K9me1, H3K4me1, H2AFZ, H3K27me3, H3K79me2, H4K20me1).

The length distribution between the negative and positive data are different. To match their lengths, we filtered out too long regions from negative data (> 2000 bp), we also dropped all regions from chromosome X (as it was done in Zhang *et al*. 2020). Additionally, we filtered out regions that overlapped with positive data regions. After filtering out sequences greater than 2000 Bps and regions with chromosome X from our negative data, we decreased the number of regions from *36 597 to 20 275*.

**Fig 2.2:**
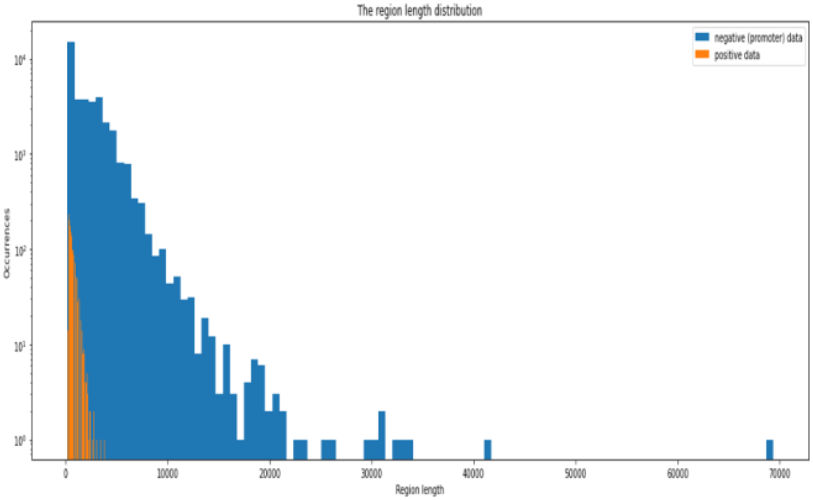
Triplex DNA site length distributions for positive & second negative (promoters) dataset for TriplexFPM-HMM

### Model Construction for Second Negative (Promoters) Dataset TriplexFPM-HMM

The amount of our negative data is almost 8 times bigger than the amount of positive data. Again, to deal with this unbalanced data, we oversampled our positive data and undersampled our negative data. We fixed the final data size to be 8000. We splitted our fixed data into three sets; *train (0*.*8), validation (0*.*1) and test (0*.*1) sets*.

### Hyperparameter Tuning for Second Negative (Promoters) Dataset TriplexFPM-HMM

We set other internal and external hyperparameters as follow: epochs (100), learning rate at (1e-3), input size (12, 1) Kernel size (32), verbose (1) and used Keras-Tensorflow to train our models. We used six different metrices in assessing the performance of our models, these include: binary accuracy, precision, recall and AUC, F1-score, MCC. We also used ReLU activation as our hidden layer activation function and Sigmoid activation function as our output layer activation function for both MLP and CNN. We used Adam optimization algorithm as our optimizer and since it is a binary classification task, we also used binary cross entropy. We used different kernel sizes with different learning rates since it increases the performance of the models. We employed early stopping criteria on the optimization procedure when the values of the loss function on the validation set stopped decreasing.

## Results and discussion

In addition to sequence-based feature models published as “Predicting RNA:DNA Triplex Structures from Sequence Features Using Deep Learning Architectures”, we developed four models incorporating histone modification marks as input features (TriplexFPM-HMM) to evaluate their predictive potential for RNA:DNA triplex formation. These models were trained using two sets of negative DNA site data: (i) generic negative data (DNA1) and (ii) promoter regions (DNA2). The results (Table 1.1) indicate that while the initial performance of these models was relatively modest compared to the lncRNA/DNA sequence-based TriplexFPM, grid search optimization substantially improved their predictive accuracy.

**Table 1.1:** TriplexFPM-HMM (Histone Modification Mark–based models)

| Model | Negative Data Source | Feature Type | Mean AUC (Before Grid Search) | Mean AUC (After Grid Search) | Kernel Size | Learning Rate |
| --- | --- | --- | --- | --- | --- | --- |
| DNA1_MLP | Negative Data 1 | Histone Marks | 0.58 | 0.59 | 32 | 1e-2 |
| DNA1_CNN | Negative Data 1 | Histone Marks | 0.51 | 0.56 | 32 | 1e-2 |
| DNA2_MLP | Negative Data 2 (Prom.) | Histone Marks | 0.67 | 0.76 | 32 | 1e-3 |
| DNA2_CNN | Negative Data 2 (Prom.) | Histone Marks | 0.57 | 0.78 | 32 | 1e-3 |

For the DNA1 set, **DNA1_MLP** initially achieved the highest mean AUC of **0.58** at a kernel size of 32 and learning rate of 1e-2, while **DNA1_CNN** performed lowest at **0.51**. Following hyperparameter tuning with grid search, performance improved slightly, with **DNA1_MLP** reaching **0.59** and **DNA1_CNN** improving to **0.56**. Although these gains were modest, they demonstrate the sensitivity of histone-based models to hyperparameter optimization.

**Fig 3.1:**
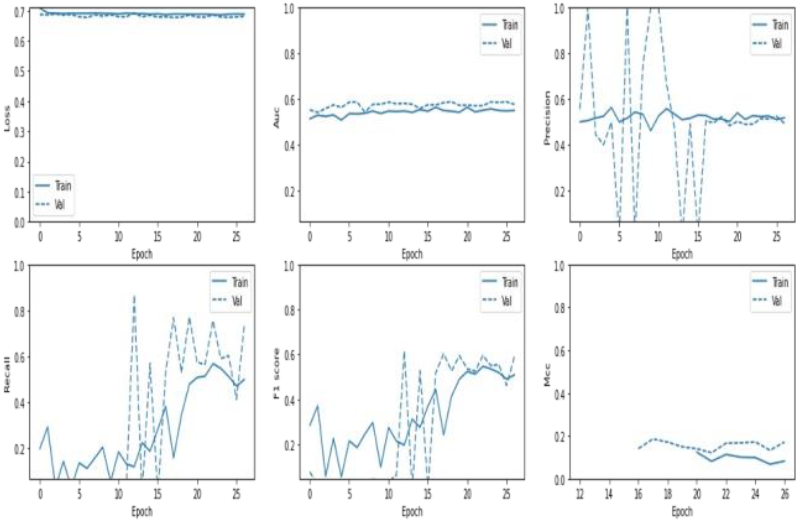
MLP model performance plot for first negative DNA site data for TriplexFPM-HMM

**Fig 3.2:**
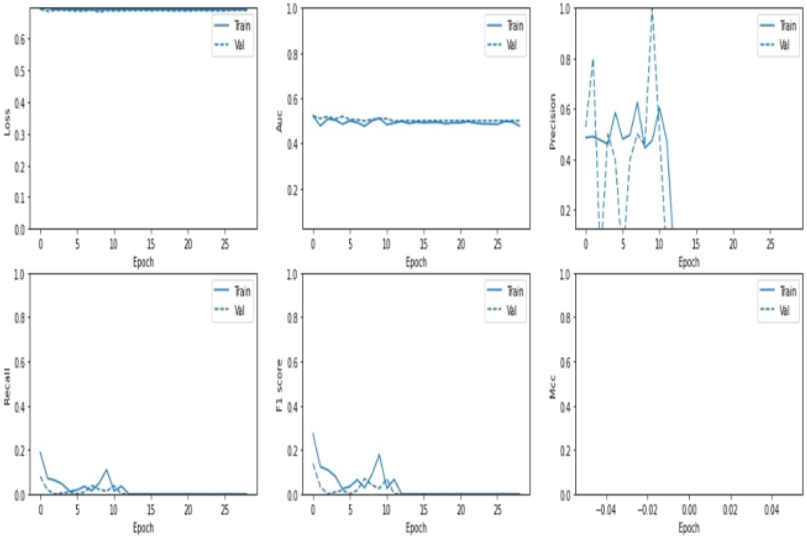
CNN model performance plot for first negative DNA site data for TriplexFPM-HMM

**Fig 3.3:**
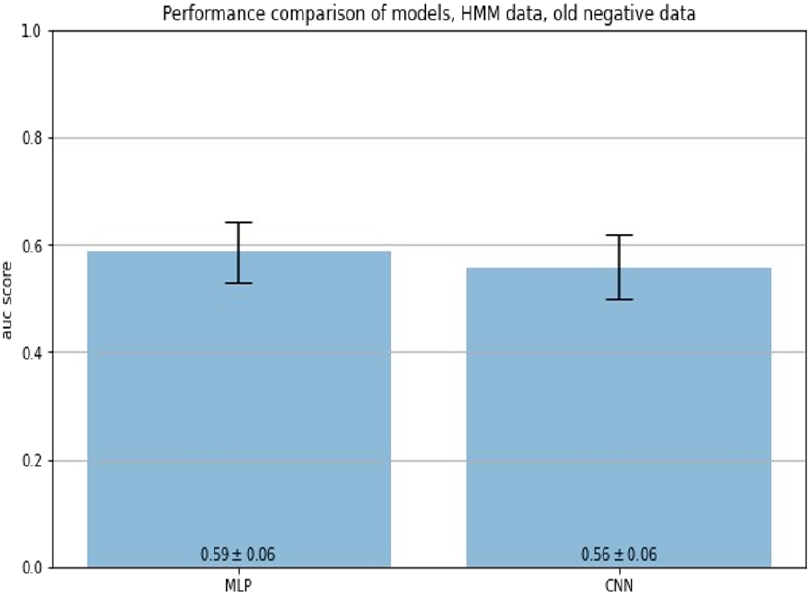
Performance comparison of MLP & CNN TriplexFPM-HMM on first negative DNA site data *after* grid search on mean AUC

For the DNA2 promoter set, performance was stronger overall. Before optimization, **DNA2_MLP** achieved a mean AUC of **0.67**, outperforming **DNA2_CNN** at **0.57**. After grid search optimization, both models showed substantial improvement: **DNA2_CNN** outperformed all histone-based models with a mean AUC of **0.78**, while **DNA2_MLP** increased to **0.76**. These findings indicate that promoter regions with histone modification marks carry stronger signals for predicting triplex formation compared to more generic DNA negative sets.

**Fig 3.4:**
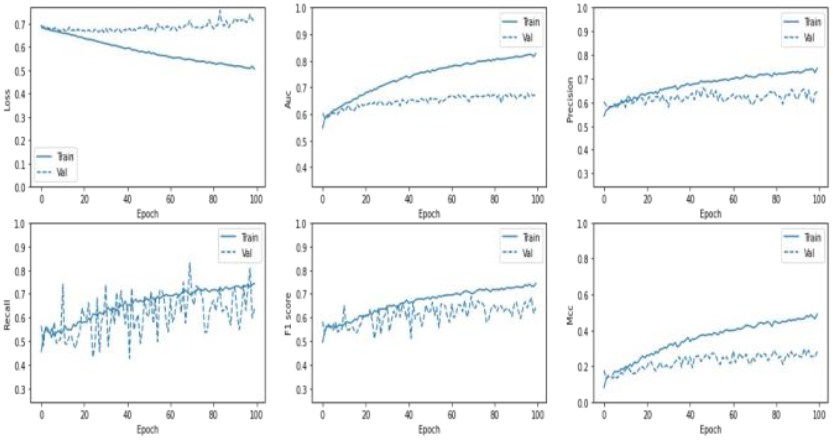
MLP model performance plot for second negative DNA site (promoters) data for TriplexFPM-HMM

**Fig 3.5:**
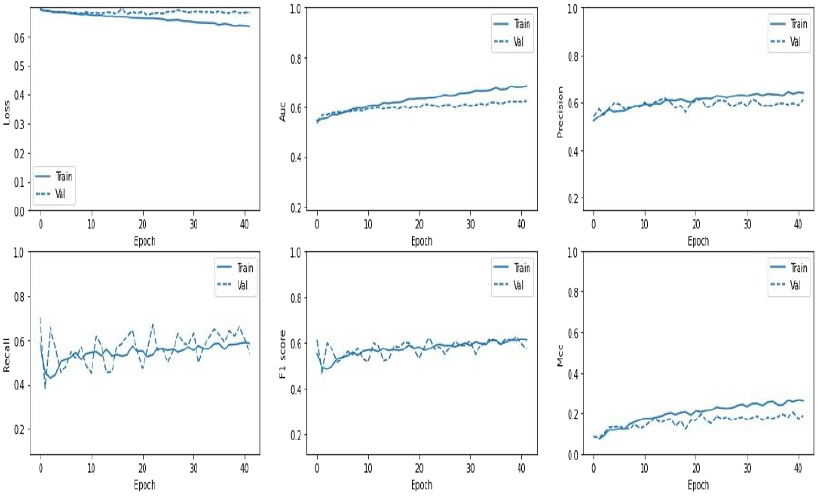
CNN model performance plot for second negative DNA site (promoters) data for TriplexFPM-HMM

**Fig 3.6:**
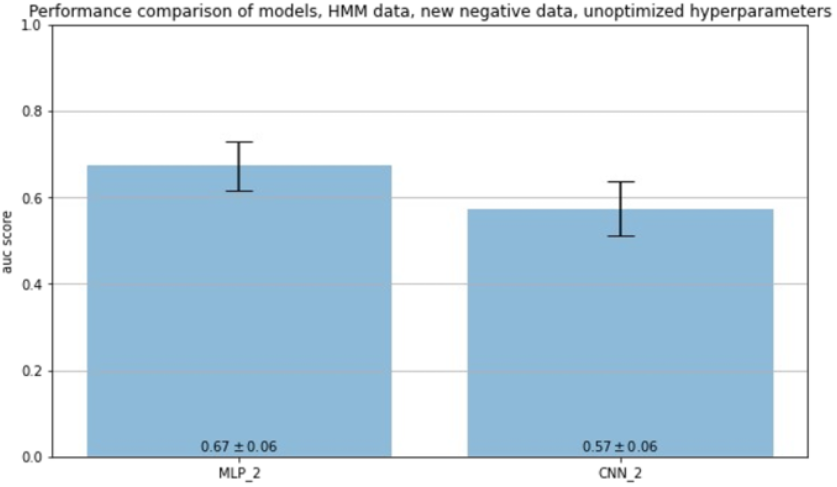
Performance comparison of MLP & CNN TriplexFPM-HMM on second negative DNA site (promoters) data *before* grid search on mean AUC

**Fig 3.7:**
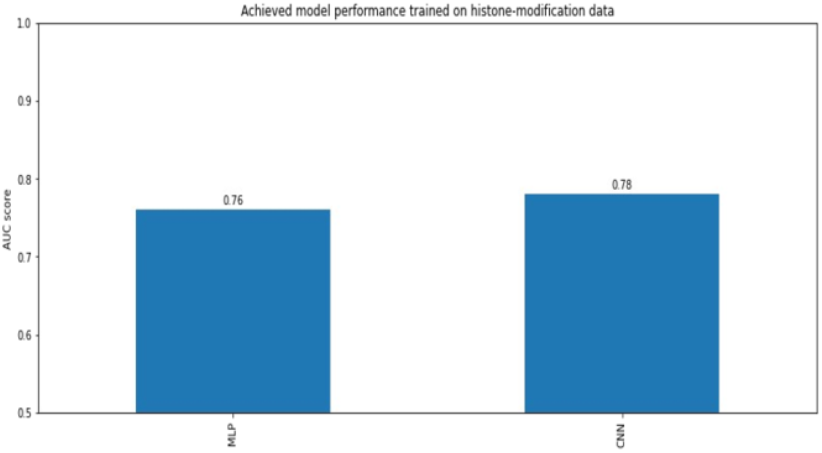
Performance comparison of MLP & CNN TriplexFPM-HMM on second negative DNA site (promoters) data *after* grid search on AUC

Collectively, these results suggest that while histone modification features alone may not achieve the same predictive power as lncRNA/DNA sequence-based models, they nonetheless provide complementary information that can enhance triplex prediction. In particular, the superior performance of promoter-based models aligns with previous findings that chromatin states, such as H3K4me3 and H3K36me3 marks, are associated with lncRNA transcriptional regulation and may predispose certain DNA regions to triplex formation.

### Limitations

Despite the improvements obtained through optimization, the overall performance of TriplexFPM-HMM remained lower than TriplexFPM sequence-based models in our previous paper. This may be attributed to the complexity of chromatin signatures, the limited size of our negative datasets, and the fact that histone modification marks were used as the sole input features without integration of additional sequence-level information.

### Future Directions

Future work should aim to integrate histone modification marks with lncRNA and DNA sequence features in multimodal architectures, potentially leveraging attention-based mechanisms to capture interactions across feature types. Moreover, larger and more diverse histone datasets are needed to increase model robustness. Comparing the performance of deep learning models with classical machine learning approaches such as support vector machines and random forests may also clarify whether simpler algorithms can capture the predictive signal in histone features. Finally, experimental validation remains essential to verify whether the histone modification marks identified by these models indeed correspond to biologically relevant triplex-forming sites.

## Declarations

### Ethics Approval and Consent to Participate

Not applicable

### Consent for Publication

Not applicable

### Availability of Data and Materials

The datasets and deep learning models used in this study are openly available on GitHub at https://github.com/Joseph-Luper-Tsenum/Using-Deep-Learning-with-Different-Architectures-to-Recognize-RNA-DNA-Triplex-Structures-from-HMMs/tree/main

Our previously published TriplexFPM sequence-based models can be found on GitHub: https://github.com/Joseph-Luper-Tsenum/Predicting-RNA-DNA-Triplex-Structures-from-Sequence-Features-Using-Deep-Learning-Architectures/tree/main

### Conflicts of interest

The author and contributors declare no conflict of interests for this article.

### Funding

This research was funded by the federal budget within the quota of the Russian Government, 2019–2021.

### Authors*’* Contribution

JLT designed the study, performed the analyses, and wrote the manuscript.

## Acknowledgments

The authors thank the Open Doors Olympiad for providing the scholarship, the Russian Government for funding, Phystech School of Biological and Medical Physics (FBMP) at the Moscow Institute of Physics and Technology (MIPT) for their support, and the members of the International Laboratory of Bioinformatics at HSE University for their valuable assistance and guidance.

## Abbreviations

CNN: Convolutional neural network
RNN: Recurrent neural network
ResNN: Residual neural network
LSTM: Long short-term memory
MLP: Multilayer perceptron
RNA: Ribonucleic acid
DNA: Deoxyribonucleic acid
lncRNA: long non-coding RNA
TriplexFPM: Triplex Forming Prediction Model
AUC: Area under the Receiver operating characteristic (ROC) Curve
ReLU: Rectified Linear Unit
lr: Learning rate
MCC: Matthews correlation coefficient
k-mer: A k-mer is a subsequence of length k derived from a longer nucleotide or amino acid sequence.

## Notes

https://github.com/Joseph-Luper-Tsenum/Using-Deep-Learning-with-Different-Architectures-to-Recognize-RNA-DNA-Triplex-Structures-from-HMMs/tree/main

### Competing Interest Statement

The authors have declared no competing interest.

### Summary of Updates

This version of the manuscript has been revised to update authorship.

